# Brain Age Prediction: Deep Models Need a Hand to Generalize

**DOI:** 10.1101/2024.06.11.598576

**Authors:** Reza Rajabli, Mahdie Soltaninejad, Vladimir S. Fonov, Danilo Bzdok, D. Louis Collins

**Affiliations:** McConnell Brain Imaging Centre, Montréal Neurological Institute, McGill University, Canada H3A 2B4; Mila - Quebec Artificial Intelligence Institute, Canada H2S 3H1

**Keywords:** Structural MRI, Robust Preprocessing Pipeline, Convolutional Neural Network, Out-of-sample Performance, Reproducibility, Reliability

## Abstract

Predicting brain age from T1-weighted MRI is a promising marker for understanding brain aging and its associated conditions. While deep learning models have shown success in reducing the Mean Absolute Error (MAE) of predicted brain age, concerns about robust and accurate generalization in new data limit their clinical applicability. The large number of trainable parameters, combined with limited medical imaging training data, contribute to this challenge, often resulting in a generalization gap where there is a significant discrepancy between model performance on training data versus unseen data.

In this study, we assess a deep model, SFCN-reg, based on the VGG-16 architecture, and address the generalization gap through comprehensive preprocessing, extensive data augmentation, and model regularization. Using training data from the UK Biobank, we demonstrate substantial improvements in model performance. Specifically, our approach reduces the generalization MAE by 44% (from 5.25 to 2.96 years) in the Alzheimer’s Disease Neuroimaging Initiative dataset and by 22% (from 4.35 to 3.40 years) in the Australian Imaging, Biomarker and Lifestyle dataset. Furthermore, we achieve a 29% reduction in scan-rescan error (from 0.86 to 0.61 years) while enhancing the model’s robustness to registration errors. Feature importance maps highlight anatomical regions used to predict age.

These results highlight the critical role of high-quality preprocessing and robust training techniques in improving accuracy and narrowing the generalization gap, both necessary steps towards the clinical use of brain age prediction models. Our study makes valuable contributions to neuroimaging research by offering a potential pathway to improve the clinical applicability of deep learning models.

## Introduction

Estimating brain age solely from T1-weighted (T1w) Magnetic Resonance Imaging (MRI) offers insight into the similarity between an individual subject’s brain and a cohort of healthy brains of various ages. Comparing the subject’s chronological age with a model-estimated brain age can represent a deviation from typical healthy aging patterns and can be used as a feature to check whether the brain is undergoing accelerated aging or not and may serve as an indicator of progression in neurodegeneration diseases. Numerous studies have established correlations between the *brain age gap* — defined as the difference between estimated brain age and chronological age — and various stages of neurodegenerative diseases. Cole et al. demonstrated an increase in apparent brain age in HIV-positive adults [1]. He and his colleagues also found an association between elevated brain age and a higher risk of death [2]. Bøstrand et al. showed that alcohol use can accelerate brain aging [3]. Numerous studies have investigated the relationship between accelerated brain aging and dementia. For instance, Wang et al. observed a significant association between brain age gap and dementia risk [4]. Biondo et al. suggested that brain age could aid in the early detection of dementia [5]. Our group found that the brain age gap significantly correlates with 18F-NAV-4694 and 18F-MK-6240 Standardized Uptake Value Ratios (SUVRs), which are tracers for amyloid-beta and tau tangles, respectively. Additionally, Beck et al. demonstrated that brain age gap is associated with cardiometabolic risk factors [6].

Due to their ability to automatically extract intensity-based, shape-related, and textural features, deep learning models have emerged as the preferred choice for brain age prediction. However, training deep learning models for medical imaging data is challenging due to limited amounts of available data, inadequate representation of the overall population, and sparsity and potential label imbalances for training and testing. In addition, many researchers fail to evaluate their models on independent datasets that have not been seen previously during training or testing. Furthermore, out-of-distribution testing is very rare as most report only cross-validation results [7]. This oversight makes it increasingly difficult to ascertain the generalizability of published models for use in other studies. Since machine learning models can be misled by inter-site MRI acquisition differences (as demonstrated by Glocker et al. [8]), the resulting lack of generalizability obstructs effective deployment of a previously trained model outside its original context. This weakness also prevents the extraction of potentially biologically-plausible interpretation of the data, hindering our understanding of underlying disease mechanisms and other biological processes. Therefore, it is crucial to exercise extra caution when (1) training a model and (2) to rigorously evaluate its performance on an independent, previously unseen dataset using various metrics to ensure robust generalization.

## Background

By searching PubMed using the keywords “MRI,” “Age,” and “Prediction,” we identified a total of *155* papers published between *2020* and *2024*. Looking more closely, we found *40* relevant papers that (1) trained a deep model on T1w MRIs to predict brain age and (2) reported results from at least one validation strategy. While successful to varying degrees, they all potentially have at least one of the three following problems: either 1) no independent data is used to assess how well the models generalize when facing out-of-distribution samples, or 2) there is a significant drop in accuracy when testing the models on external data, or 3) no results are reported regarding the model’s robustness, such as how the output changes with small perturbations to the input, like registration errors or scan-dependent differences.

We review a number of these papers here in chronological order. Table 1 summarizes our findings. Feng et al. [9] trained a 3D Convolutional Neural Network (CNN) with multiple interleaved convolutional blocks to predict brain age. They tested their model on an independent dataset, Cam-CAN, and reported a Mean Absolute Error (MAE) of 4.21 years for predicting an individual’s age from T1w MRI structural scans. These authors also ran a reproducibility test and showed that their model prediction remains consistent across sessions, with an approximate standard deviation of 1 year.

**Table 1.** Deep brain age prediction papers discussed in the introduction and tests they reported. Test #1 is out-of-distribution test and Test #2 is robustness test. The sign “✓✓” indicates no problem, the sign “✓” indicates some tests were done, but there are possible shortcomings, and the sign “✗” indicates no proper testing was done/reported.

| Paper | Test #1 | Test #2 | Comments |
| --- | --- | --- | --- |
| Feng et al. [9] | √ | √ | Independent testing was done on only one dataset. Also, just one reproducibility test is reported which was done only on 3 subjects. |
| Lombardi et al. [10] | X | X | Hold-out data came from the same distribution with training data. |
| Kuo et al. [11] | X | X |  |
| Dinsdale et al. [12] | X | X |  |
| Ren et al. [13] | X | X |  |
| Besson et al. [14] | X | X |  |
| Hepp et al. [15] | X | X |  |
| Peng et al. [16] | X | X |  |
| Leonardsen et al. [17] | √√ | X |  |
| Lee et al. [18] | √ | X | Independent testing was done on only one dataset. |
| Dular et al. [19] | √ | √ | Independent testing was done only on the UKBB dataset, leading to potentially more optimistic results. They did not perform any robustness test for errors related to the preprocessing pipeline. |

Lombardi et al. [10] trained a deep model on morphological features extracted by FreeSurfer v5.3.0 and tested their model on previously unseen samples from previously seen scan sites. They reported an MAE of 2.7 years on the test set. Additionally, they did not study the impact of possible errors in different stages of the FreeSurfer *recon-all* processing command. Kuo et al. [11] reported an MAE of 3.33 years for predicting brain age of the PAC2019 dataset subjects, but no independent testing and no sensitivity/stability analyses were performed.

Dinsdale et al. [12] reported MAE that ranged from 2.71 to 3.09 years. However, they did not test their model on unseen images from a dataset different from the training dataset. Also, despite training three networks simultaneously, suggesting potential increased robustness, they did not conduct any specific robustness evaluation.

Ren et al. [13] achieved an MAE of 2.65 years as their best reported result, however they did not use unseen external data to test their deep model. Additionally, although they used registered MRIs as training data, they did not report how sensitive their model is when faced with different sources of variabilities such as registration errors or scanner-related artifacts.

Besson et al. [14] reduced the number of trainable parameters of their model by exploiting a Graph CNN architecture and achieved test MAEs of 2.73 to 8.61 years on different datasets. While they tested their model on hold-out data, the testing data came from the same datasets as that used for training, thus came from the same distribution. Moreover, although they registered all images to the Montreal Neurological Institute (MNI) space and extracted surface vertices, they did not report any robustness testing.

Hepp et al. [15] introduced an uncertainty measure to have an estimation of the predicted brain possible age range, instead of having only a single value as the predicted brain age. They reported an MAE of 3.21 ± 2.45 years. They trained and tested their model on the same dataset, the Germany National Cohort (GNC) study. No external or robustness test was done.

Peng et al. [16] employed preprocessed MRIs. They reduced the number of trainable parameters significantly by removing the fully-connected layer from their model architecture and achieved an MAE of 2.18 years. However, they did not validate their model externally, nor did they test their model robustness. Leonardsen et al. [17] modified Peng’s model and trained it on ∼53K MRIs from multiple studies and reported an MAE of 2.47 years on the internal (within domain) test set and 3.90 years on average on external (out of domain) test images from other studies. This 1.43 year increase in MAE is indicative of the generalizability limit of their model in out of domain data. They did not test the robustness of their model.

Lee et al. [18] trained a model on ∼2K MRIs from the Mayo dataset [20] and tested it on Alzheimer’s Disease Neuroimaging Initiative (ADNI) [21] subjects. They concluded that their model is generalizable since the internal (within domain) and the external (out of domain) test accuracies were statistically comparable (MAE of 3.48 years and 3.51 years respectively after bias correction). While their model was trained on registered MRIs, they did not report on the impact of small input disturbances on the prediction accuracy, nor did they mention any use of any method to make their model more robust.

Recently, Dular et al. [19] trained 4 different models on ∼2.5K T1w MRIs gathered from multiple sites. They did two external validations: first, testing the model on samples from an unseen site (MAE: 3.31 years for within domain to 3.65 years for out of domain testing for different models and different offset correction strategies), and second, testing the model on samples that had gone through a different preprocessing pipeline (MAE: 3.71 to 9.80 years).

The study revealed that the models’ error is significantly linked to the preprocessing pipeline. While they include different tests in their framework to assess the robustness of the trained models, they did not test those models’ robustness in the face of preprocessing pipeline-related errors (e.g. registration errors). It is also noteworthy that their external testing relied exclusively on data from the UK Biobank (UKBB) dataset [22], widely acknowledged as one of the highest-quality, homogeneous datasets available in the field. This choice of dataset may potentially lead to more optimistic test results.

## Materials and methods

### Datasets

The T1w MRI data from five datasets used to train, validate, and test our Brain Age deep learning model are described below.

### UK Biobank (UKBB)

The UKBB is an on-going project with the aim of gathering a large dataset of biomarkers to study human health. We downloaded version v14940, containing 39,676 T1w MR images of healthy subjects. All images were scanned using the same scanner, a Siemens Skyra 3T, with a 3D MPRAGE sequence (TR = 2000 ms, TE = 2.01 ms, TI = 880 ms, and flip angle = 8°) with 1 mm x 1 mm x 1 mm voxels.

### Alzheimer’s Disease Neuroimaging Initiative (ADNI)

ADNI was launched in 2003 as a public-private partnership, led by Micheal W. Weiner, MD, with the primary aim of investigating whether MRI, Positron Emission Tomography (PET), other biological markers, and clinical and neuropsychological assessment can be combined to measure the progression of AD. In this study we selected baseline scans of healthy participants from ADNI-1, ADNI-2, and ADNI-Go cohorts consisting of 594 T1w MRI scans of 280 cognitively normal subjects at baseline (first visit) acquired with an MPRAGE sequence (TR = 2300/2400 - 3000 ms, TE = ∼3 ms, TI = 900 - 1000 ms, and flip angle = 8 - 9°). The images were scanned using different scanners (from 3 different manufacturers: GE, Siemens, and Philips) at different sites. The resolution of images in ADNI-1 is 0.9375 mm x 0.9375 mm x 1.2 mm and in ADNI-2/Go is 1 mm x 1 mm x 1.2 mm [23].

### Australian Imaging Biomarkers and Lifestyle Study of Aging (AIBL)

AIBL is a study to discover which biomarkers, cognitive characteristics, or life-style factors can lead to AD development. From this study we selected baseline scans of cognitively normal participants, consisting of 245 T1w MR images scanned on a Siemens Avanto 1.5T with an MPRAGE sequence (TR = 1900 ms, TE = 2.13 ms, TI = 900 ms, flip angle = 9°). The resolution of scans is 1 mm x 1 mm x 1.2 mm. Only baseline images were used.

### Open Access Series of Imaging Studies 3 (OASIS-3)

OASIS is an open-access compilation of data for more than 1,000 subjects collected from different on-going projects over 30 years. All OASIS-3 scans collected on Siemens TIM Trio 3T using MPRAGE sequence (TR = 2400 ms, TE = 3.08 ms, TI = 1000 ms, flip angle = 8°). The resolution is 1 mm x 1 mm x 1 mm. In this study, we selected 859 baseline scans from cognitively normal participants.

### Movement-related artifacts (MR-ART)

MR-ART is an open-access dataset of structural brain MRIs collected from 148 cognitively normal subjects including motion-free and motion-affected data acquired from the same participants in the same session with the goal of raising awareness about the impact of motion on MRI-derived data and evaluating approaches to correct for them. All scans were collected on a Siemens Magnetom Prisma 3T using an MPRAGE sequence (TR = 2300 ms, TE = 3 ms, TI = 900 ms, flip angle = 9°). The resolution is 1 mm x 1 mm x 1 mm.

The UKBB images were employed for training our models, while testing was conducted on four other datasets. We specifically opted for baseline scans (those taken during the first visit) from each dataset, with an exception for datasets lacking a longitudinal structure to ensure the robustness of our model evaluation and mitigate potential challenges arising from correlations between different test samples.

### Data availability

OASIS3 and MR-ART are open-access. UKBB, ADNI, and AIBL are available upon request and approval.

### Preprocessing

All T1w MRI data used in this study (including training, evaluation, and test phases) were preprocessed using the following steps: 1) brain extraction using SynthStrip [24], 2) intensity normalization through histogram matching [25], 3) denoising with adaptive non-local means [26], 4) N4 bias field correction [27], 5) repeating step 2, and 6) affine registration to the MNI152 nonlinear [28], [29] symmetric template using ANTs [30]. The size of the output image was 192×192×192 with 1 mm^3^ voxels. These volumes were used for all model fitting, validation, and testing. For each preprocessing step, we used some of the most advanced and reliable methods available in the literature, while many other studies rely on older approaches or utilize traditional FreeSurfer recon-all pipelines, which incorporate older methods such as the skull stripping technique introduced by Ségonne et al. [31] and the N3 bias field correction [32]. Additionally, we performed intensity normalization twice during the pipeline: once before bias field correction and once after.

### Quality Control

We manually inspected all preprocessed images using Qrater [33]. Approximately ∼4% of the preprocessed images failed, mostly due to either corrupted brain masks or failed registration. The number of failed images in each dataset are as follows: 1,976 of UKBB, 47 of ADNI, 1 of AIBL, and 58 of OASIS-3. The demographics of the subjects that passed the QC are summarized in Table 2.

**Table 2.** Demographic information of the various datasets utilized in this study.

| **Dataset** | **Subset** | **Size**<br>**[Subjects / Images]** | **Sex**<br>**[F / M]** | **Age Range**<br>**[min - max, mean±std]** |
| --- | --- | --- | --- | --- |
| UK BioBank (UKBB) | Full | 37,700 / 37,700 | 20299 / 17401 | 45.0 - 83.0, 64.02±7.54 |
| Alzheimer's Disease<br>Neuroimaging Initiative<br>(ADNI) | Healthy<br>Controls | 280 / 547<br>(333 1.5T, 214 3T) | 144 / 136 | 59.9 - 90.1, 74.75±5.75 |
| Australian Imaging<br>Biomarkers and Lifestyle<br>Study of Aging (AIBL) | Healthy<br>Controls | 244 / 244 | 140 / 104 | 60.0 - 92.0, 73.50 ± 6.48 |
| Open Access Series of<br>Imaging Studies 3 (OASIS-3) | Healthy<br>Controls | 611 / 801 | 258 / 170<br>(183<br>unidentified) | 42.7 - 95.3, 67.58±8.23 |
| Movement-related artefacts<br>(MR-ART) | Full | 148 / 436 | 95 / 53 | 18.2 - 74.8, 30.01±12.72 |

### Training Data

We split the UKBB data into two sets: one for training and validation (n = 95% of 37,700 = 35,815) and one for hold-out testing (n = 5% of 37,700 - 35,815 = 1,885). The training and validation set was split into 3 parts: 32,045 MRIs for training, 1,885 for validation set 1 and 1,885 for validation set 2. The two external validation sets were used to better check for any potential bias when estimating the within-domain accuracy. All sets were constructed to have an approximately uniform age distribution.

### Deep Learning Model for Brain Age Prediction

We trained three different models with the training data, which was preprocessed, and quality controlled earlier (Fig. 1). The first model is an extension of the SFCN model of Peng, called SFCN-reg [17], with identical hyperparameter choices as specified in [17]. SFCN-reg is a convolutional neural network based on the VGG16 architecture, with the exclusion of the linear layer before VGG16’s final layer, effectively reducing the number of trainable parameters to ∼3 millions. No hyperparameter tuning or additional augmentation techniques beyond those in the original paper were employed, allowing for a direct comparison of the impact of our more robust preprocessing pipeline. Fig. 1 shows the architecture of SFCN-reg used in this study.

**Figure 1.**
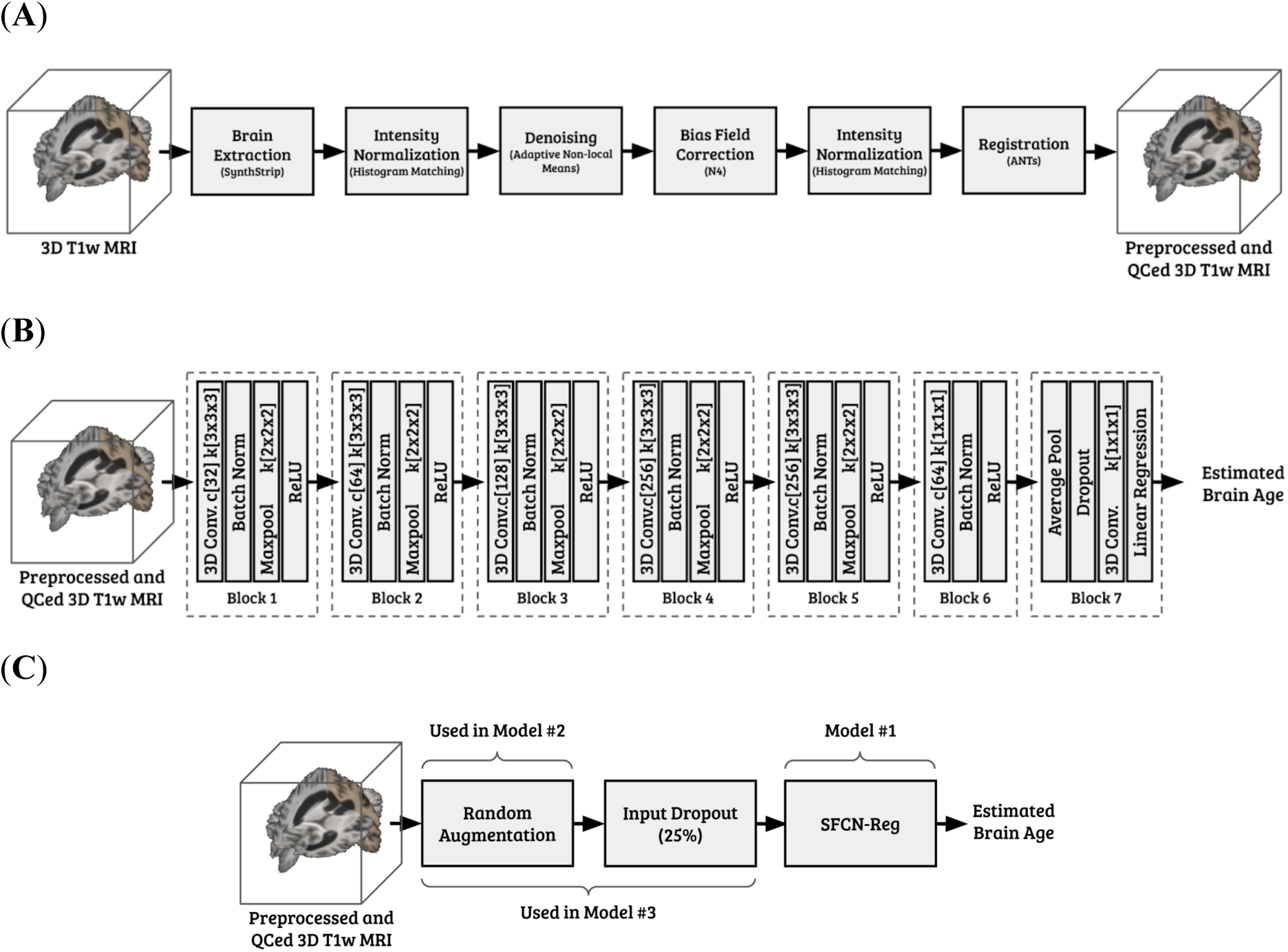
Details of the models we trained. (**A**) Our preprocessing pipeline, (**B**) The architecture of SFCN-reg we utilized, (**C**) The pipeline for the second and the third model we trained.

The second model was the same as the first, but during training we randomly applied a combination of 6 different data augmentation methods (*RandomAffine*, *RandomGamma*, *RandomBiasField*, *RandomMotion*, *RandomBlur*, and *RandomNoise* - all implemented in TorchIO library [34]) to each image before introducing it to the network (Fig. 1C). The parameters for data augmentation are described below. We hypothesized that these manipulations would make the SFCN-reg network more robust to data variability seen in the within domain training data and eventually, in the out of domain testing data.

The third model was the same as the second, but during the training before feeding an MRI to the network, we randomly masked 20% of the voxels in the image, also known as input dropout (Fig. 1C).

To finalize each model, we identified the epoch with the best average validation error from validations sets 1 and 2. We then evaluated the models with both internal (in-distribution) testing with the 5% held out independent UKBB dataset and with external (out-of-distribution) testing on ADNI, AIBL, OASIS-3 and MR-ART datasets. It is crucial to note that none of these test samples were seen by the model during the training/validation.

### Data Augmentations

We utilized the following augmentation techniques implemented in TorchIO library [34] when training the second and the third model:

- **RandomAffine:** Applies a random affine transformation (translation, rotation, and scaling) to the input. Translation parameters {*dx*, *dy*, *dz*} were randomly chosen from a continuous uniform distribution of (−2 mm, 2 mm). Rotation parameters {*rx*, *ry*, *rz*} were randomly chosen from a continuous uniform distribution of (−2°, 2°). Scaling parameters {*sx*, *sy*, *sz*} were randomly chosen from a continuous uniform distribution of (−0.99, 1.01). We selected parameters to represent ∼1% error in registration.
- **RandomGamma:** Randomly adjusts contrast values (range of [0, 1]) by raising them to the power of *ψ*. *ψ* values were calculated as *e^β^*, with *β* chosen from a continuous uniform distribution of (−0.3, 0.3). This shifts the intensity distribution of an image nonlinearly to mimic the variability due to different scanner parameters.
- **RandomBiasField:** Applies a bias field (linear combination of polynomial basis functions) to the entire image. Although we used the N4 method to remove non-uniformity from images, complete removal of the bias field is not guaranteed. Therefore, we used a first-order polynomial function with a coefficient randomly chosen from a continuous uniform distribution of (−0.3, 0.3).
- **RandomMotion:** Simulates several movements in the form of translation or rotation. Since we discarded images with extensive movement during the QC phase, we simulated head movement by repeating the original image 1 to 4 times (uniformly chosen). Each repetition involved translating the original image by d voxels chosen from a continuous uniform distribution of (−3, 3).
- **RandomBlur:** Applies a Gaussian filter with randomly chosen standard deviations across different axes. Based on our visual inspection, we set a standard deviation of no larger than 1.5 in each axis.
- **RandomNoise:** Applies Gaussian noise with a randomly chosen mean and standard deviation to the image. According to our visual inspection, we chose the mean 0 and a randomly chosen standard deviation from uniform distribution of (0, 0.06) for a signal range of [0, 1].

See Fig. 2, for the impact of each augmentation method on a sample image from UKBB.

**Figure 2.**
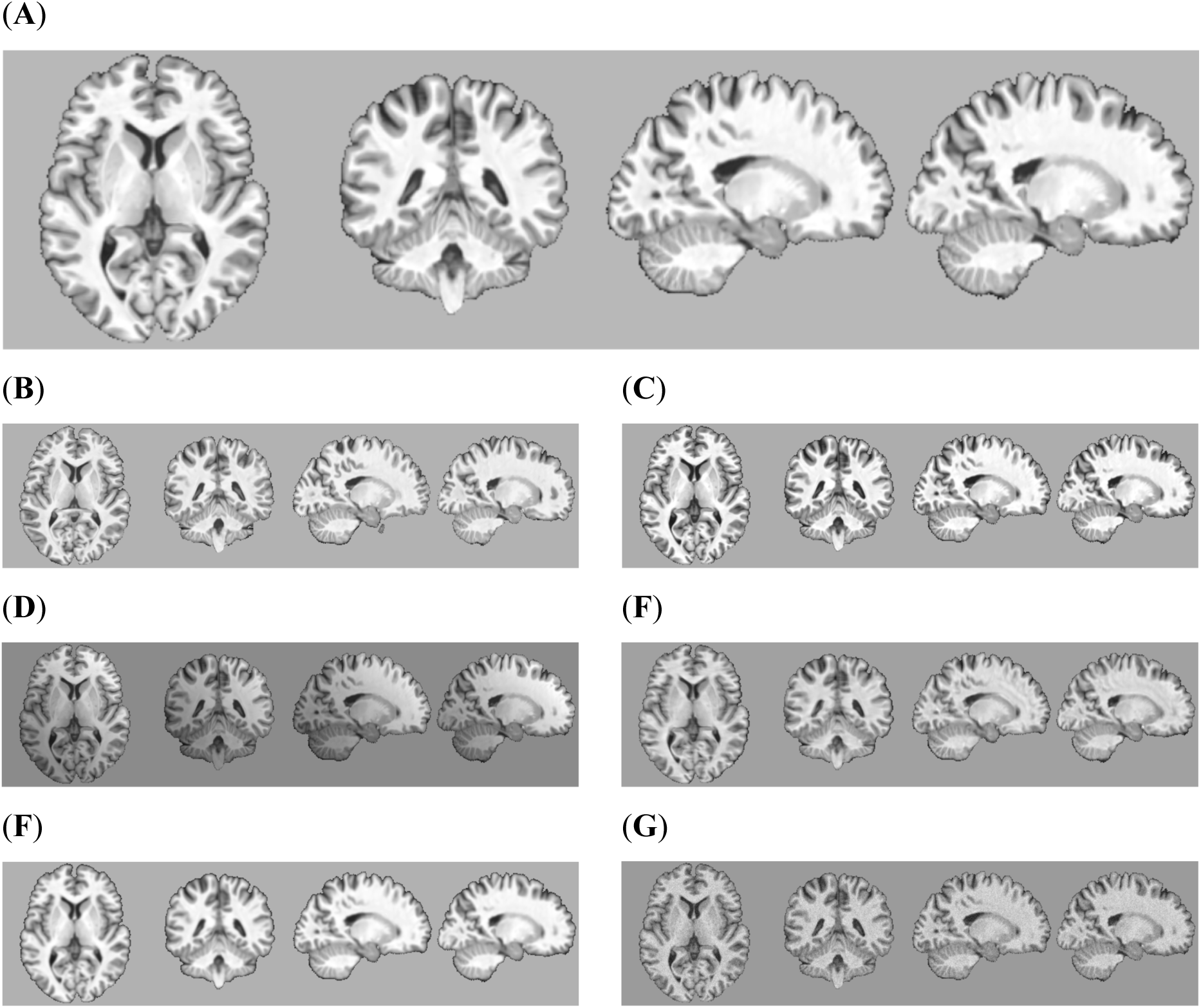
The impact of each TorchIO augmentation method on a sample image from UKBB. (**A**) The original sample. (**B**) *RandomAffine* with (*dx*, *dy*, *dz*) = (−2, −1, 0), (*rx*, *ry*, rz) = (2, 1, 0), (*sx*, *sy*, sz) = (1.005, 0.995, 1.000). (**C**) *RandomGamma* with *β* = 0.25, (**D**) *RandomBiasField* with *coefficient* = 0.25. (**E**) *RandomMotion* with 3 simulated movements with *d* = 1.5. (**F**) *RandomBlur* with *std* = 1.0. (**G**) *RandomNoise* with *mean* = 0 and *std* = 0.05

### Bias Correction

Many age prediction models have issues with *regression to the mean*, where the model predicts younger subjects as older and older subjects as younger. This bias can be observed in the correlation between prediction error and the true value. Specifically, the age prediction error tends to be more negative for older subjects on average and more positive for younger subjects. To address this bias, we applied a linear mapping to the model output to null the correlation between age prediction error and true age on the validation set (as outlined in the original SFCN paper [16]). Since correction coefficients are computed only in training, and reused with any of the test sets, there is no data leakage.

## Results

For each independent testing dataset, we performed a comprehensive set of empirical checks, including the model’s stability across different subsets of test data, the model’s ability to extrapolate, scan-rescan error assessments, and the model’s reliability in the presence of registration errors - all detailed below. This suite of evaluations aimed to provide a thorough understanding of the trained model performance across different datasets and under various conditions.

Additionally, to assess the impact of our preprocessing pipeline on achieving more generalized predictions, all test samples underwent the identical preprocessing pipeline and QC procedure (cf. above). Our hypothesis is that this standardized processing protocol safeguards against observed differences in model performance being attributed to dataset-related artifacts rather than genuine underlying factors.

### Test #1: Performance on different subsets of a test set

To assess the model’s reliability and robustness, we computed the mean and standard deviation of the Mean Absolute Error (MAE) in predicted age, in years, using a resampling procedure (like bootstrapping), selecting 50 subjects randomly from each test set 100 times. The performance of our model was compared with data published with the original SFCN-reg model in Table 3. Of note, our model was trained exclusively on UKBB data, thereby constraining the observed trained age to the range 45-81 years old. The results in Table 3 are from the full test sets, regardless of the age range in each set.

**Table 3.** Performance comparison of the original SFCN-reg and our trained models. The internal test was done on UKBB, and the external test was done on ADNI, AIBL, and OASIS3 using the full age range available (not limited only to the training age range). Mean Absolute Error (MAE) measured in years. Numbers in brackets are model performance after the regression-based bias correction. Prep: Our Preprocessing Pipeline, Aug: Data Augmentation, Drop: Input Dropout (20%).

| Dataset | UKBB | ADNI | AIBL | OASIS3 |  |
| --- | --- | --- | --- | --- | --- |
| n | 1885 | 280 | 244 | 611 |  |
| [Min, Max] | [46.0, 81.0] | [59.9, 90.1] | [60.0, 92.0] | [42.7, 95.3] |  |
| Mean ± Std | 63.9 ± 7.61 | 74.9 ± 5.75 | 73.5 ± 6.48 | 67.6 ± 8.23 | |
| Model / Test (Metric) | Internal Test (MAE) | External Test (MAE) | External Test (MAE) | External Test (MAE) | External Test (Average) |
| SFCN<sup>1</sup> | 2.18 | - | - | - | - |
| SFCN-reg<sup>2</sup> | 2.47 | 5.25 | 4.35 | - | 3.90<sup>3</sup> |
| **Model #1**<br>SFCN-reg + Prep | **2.06 ± 0.22**<br>**[2.20 ± 0.23]** | 3.44 ± 0.34<br>[2.96 ± 0.30] | 3.86 ± 0.35<br>**[3.40 ± 0.31]** | **2.93 ± 0.34**<br>**[2.84 ± 0.32]** | 3.41<br>**[3.07]** |
| **Model #2**<br>SFCN-reg + Prep + Aug | 2.31 ± 0.26<br>[2.47 ± 0.25] | **3.21 ± 0.31**<br>**[2.88 ± 0.29]** | **3.82 ± 0.35**<br>[3.44 ± 0.32] | 2.97 ± 0.38<br>[3.04 ± 0.33] | **3.33**<br>[3.12] |
| **Model #3**<br>SFCN-reg + Prep + Aug + Drop | 2.33 ± 0.24<br>[2.52 ± 0.25] | 3.38 ± 0.31<br>[3.04 ± 0.30] | 4.05 ± 0.39<br>[3.56 ± 0.36] | 3.06 ± 0.38<br>[3.05 ± 0.31] | 3.50<br>[3.22] |

<sup>1</sup> Numbers from the original paper [16]
<sup>2</sup> Numbers from the original paper [17]
<sup>3</sup> The external test average for the original SFCN-reg paper [17] includes test results on IXI and the Norwegian Cognitive NeuroGenetics Sample, and so is not directly comparable to the average of ADNI, AIBL, and OASIS.

Fig. 3A displays the scatter plots between the true chronological age and the predicted brain age from our implementation of the SFCN-reg model trained with UKBB data processed by the improved pipeline (Model #1). Internal test results in Table 3 closely match those reported in the original SFCN paper [16] and those reported in the original SFCN-reg [17], indicating the success of our SFCN-reg re-implementation. However, since our SFCN-reg implementation was trained on a narrower dataset range, a direct comparison is not possible. External test results showed up to 43% improvement on ADNI (from 5.25 years down to 3.46/3.01 years before/after bias correction), and up to 27% reduction in MAE for AIBL (from 4.35 years down to 3.52/3.19 years before/after bias correction).

**Figure 3A.**
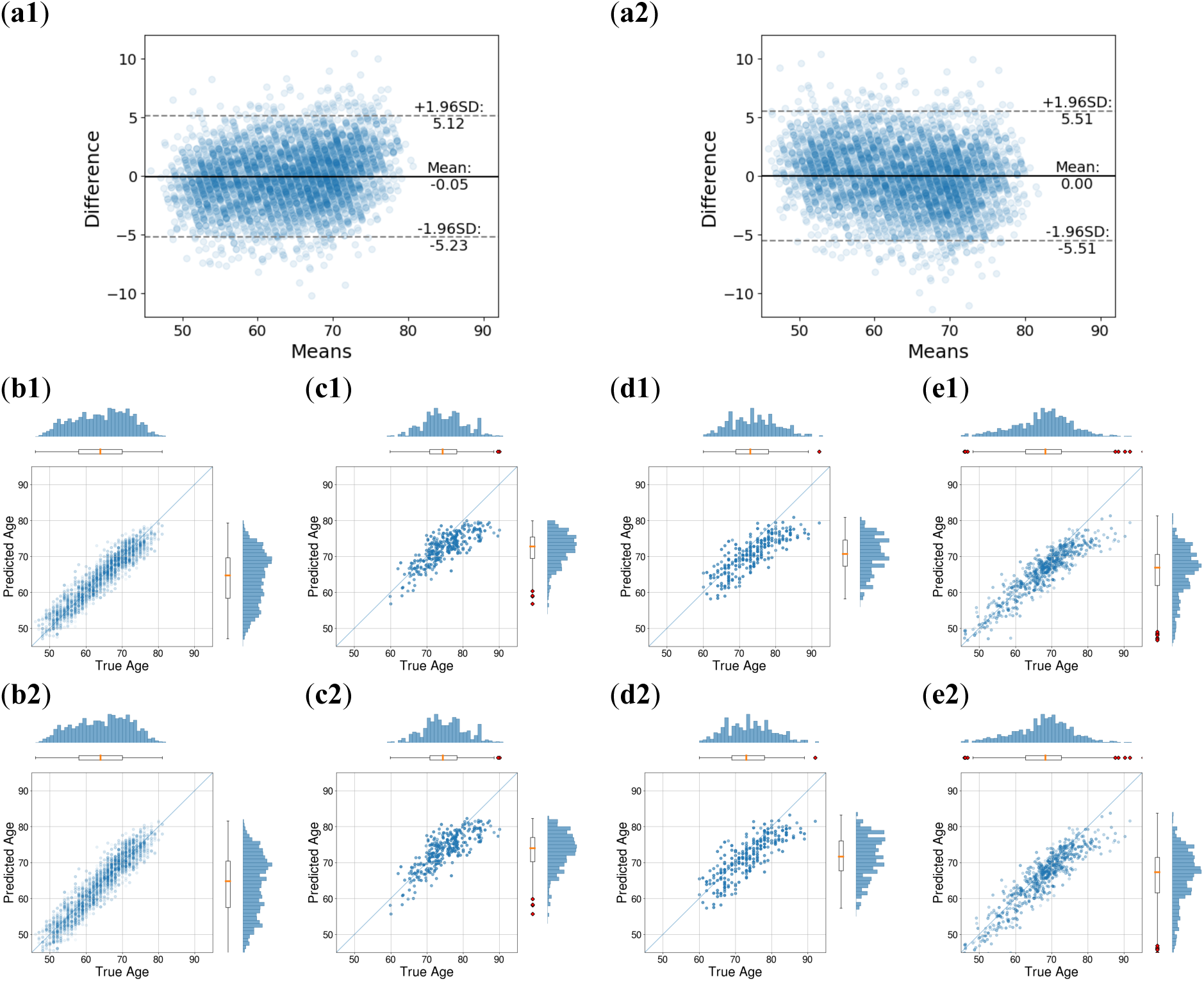
Model #1 (SFCN-reg + Preprocessing Pipeline): Predicted Age vs True (Chronological) Age. Bland-Altman plots (sub-panel (**a**)) compare model performance on the UKBB validation set. (**a1**) represents the direct deep model output, while (**a2**) shows results after regression bias correction. Additionally, sub-panel (**b**) displays the Predicted Age vs. True Age scatter plot for the UKBB validation set, with (**b1**) showing results without bias correction and (**b2**) with bias correction. Age histograms are included for both Predicted (vertical) and True Age (horizontal). Sub-panels (**c**), (**d**), and (**e**) extend the analysis to external test data from ADNI, AIBL, and OASIS, with Row 1 (**c1**, **d1**, **e1**) showing results without bias correction and Row 2 (**c2**, **d2**, **e2**) with bias correction.

Fig. 3B shows the scatter plots between the true chronological age and the predicted brain age from our implementation of the SFCN-reg model trained with UKBB data processed using our improved pipeline and with extended data augmentation during the training (Model #2). As reported in Table 3, adding data augmentation increases MAE in the internal test results (from 2.06 years to 2.31 years before bias correction and from 2.20 years to 2.47 years after bias correction, approximately 12% in both scenarios) but results in a 3% to 7% improvement of MAE for ADNI (from 3.44 years to 3.21 years before bias correction, and from 2.96 years to 2.88 years after bias correction). There is no significant change for AIBL and OASIS3 before/after bias correction.

**Figure 3B.**
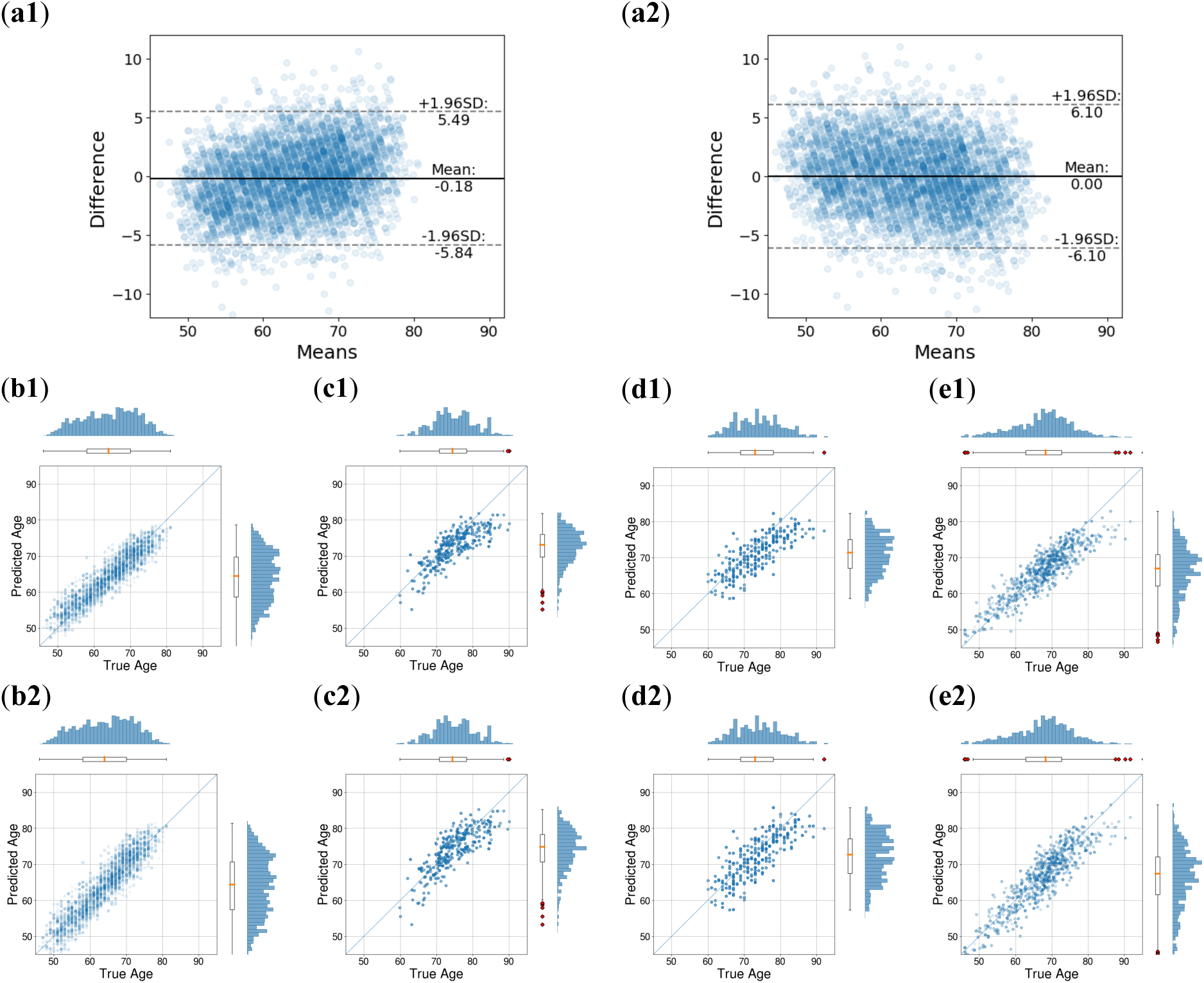
Model #2 (SFCN-reg + Preprocessing Pipeline + Augmentation): Predicted Age vs True (Chronological) Age. Bland-Altman plots (sub-panel (**a**)) compare model performance on the UKBB validation set. (**a1**) represents the direct deep model output, while (**a2**) shows results after regression bias correction. Additionally, sub-panel (**b**) displays the Predicted Age vs. True Age scatter plot for the UKBB validation set, with (**b1**) showing results without bias correction and (**b2**) with bias correction. Age histograms are included for both Predicted (vertical) and True Age (horizontal). Sub-panels (**c**), (**d**), and (**e**) extend the analysis to external test data from ADNI, AIBL, and OASIS, with Row 1 (**c1**, **d1**, **e1**) showing results without bias correction and Row 2 (**c2**, **d2**, **e2**) with bias correction.

Fig. 3C illustrates the scatter plots between the true chronological age and the predicted brain age from our implementation of the SFCN-reg model trained with UKBB data processed using our improved pipeline and with extended data augmentation during the training, as well as incorporating input dropout (Model #3). As detailed in Table 3, apart from the test results reported for ADNI, this model does not show improved results compared to the other two. Internal test results demonstrate up to 14.5% increase of MAE in different scenarios. Furthermore, there is up 6.0% increase of MAE for AIBL data, and up to 7.4% increase for OASIS3 data.

**Figure 3C.**
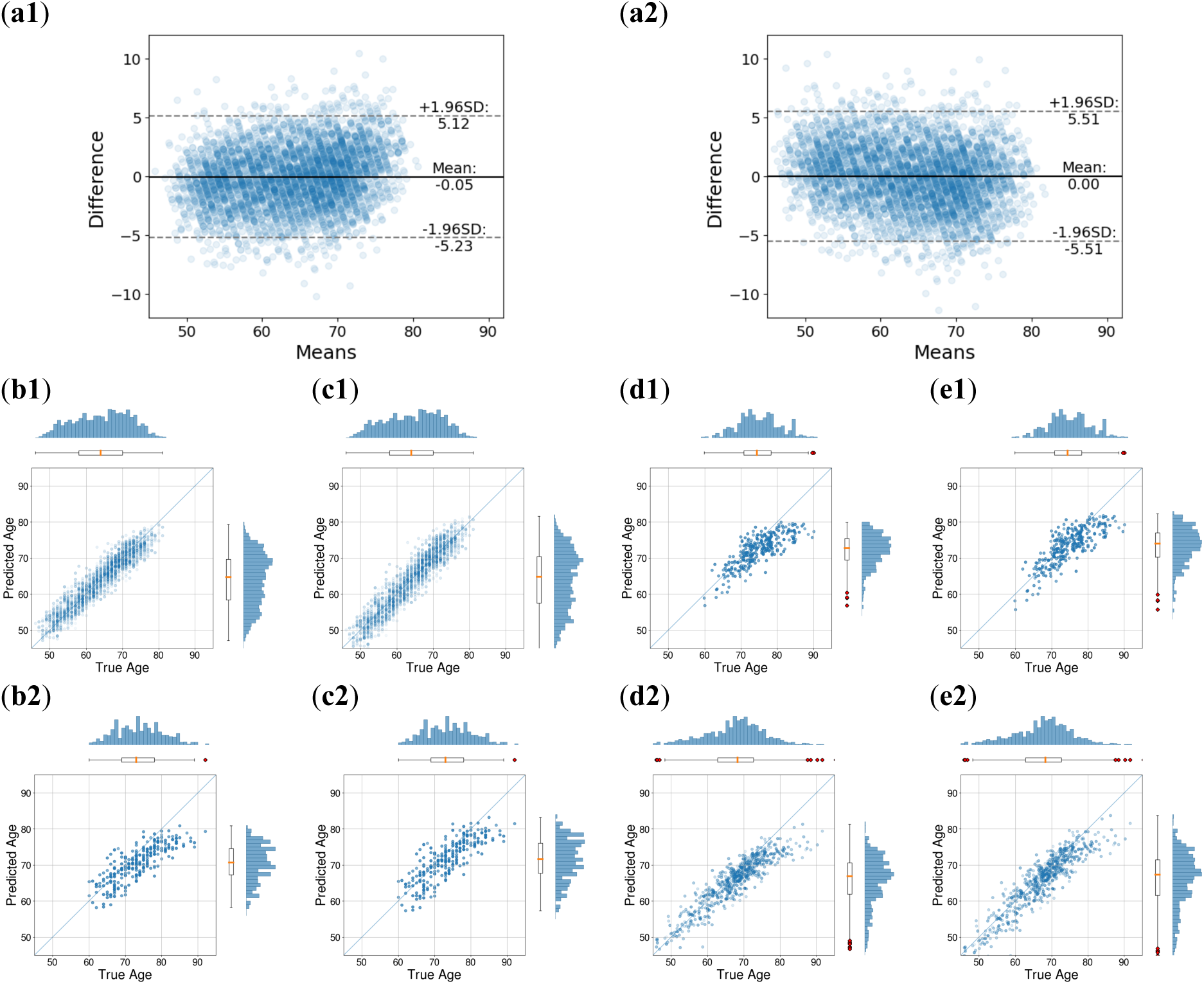
Model #3 (SFCN-reg + Preprocessing Pipeline + Augmentation + Dropout): Predicted Age vs True (Chronological) Age. Bland-Altman plots (sub-panel (**a**)) compare model performance on the UKBB validation set. (**a1**) represents the direct deep model output, while (**a2**) shows results after regression bias correction. Additionally, sub-panel (**b**) displays the Predicted Age vs. True Age scatter plot for the UKBB validation set, with (**b1**) showing results without bias correction and (**b2**) with bias correction. Age histograms are included for both Predicted (vertical) and True Age (horizontal). Sub-panels (**c**), (**d**), and (**e**) extend the analysis to external test data from ADNI, AIBL, and OASIS, with Row 1 (**c1**, **d1**, **e1**) showing results without bias correction and Row 2 (**c2**, **d2**, **e2**) with bias correction.

### Test #2: Out-of-age-range extrapolation ability

As mentioned above, all three models underwent training on a dataset comprising 32,045 samples from the UKBB, with an age range spanning from 45 to 81 years with a nearly uniform distribution. It is important to investigate the model’s ability to extrapolate to predict the brain age of individuals who fall outside the training range, a necessary characteristic to apply the model in broader demographic contexts beyond the training data. We were unable to find any previously published results that addressed the issue of extrapolation in brain age prediction for comparison. Table 4 shows the results of testing of subjects with an age greater than, or less than, that available in the training set (top half of Table 4) or that fall within the range of the training data (bottom half of Table 4). For those out of the range, most *predicted* brain ages fall within the training age range, resulting in MAEs substantially greater than observed in the previous set of experiments (see Fig. 4 for scatter plots for Model #3). Additionally, the sub-cohorts of cognitively normal subjects aged over 83 years old in ADNI, AIBL, and OASIS3 are relatively small, which complicates drawing robust conclusions. For MR-ART dataset, where the age range and the number of samples available are more than twice than other datasets, MAEs and R2 scores for different models are not conclusively different from each other. Not surprisingly, for the sub-cohort from each external test dataset that is *within* the training age range, the MAE is smaller than that reported in Table 3.

**Figure 4.**
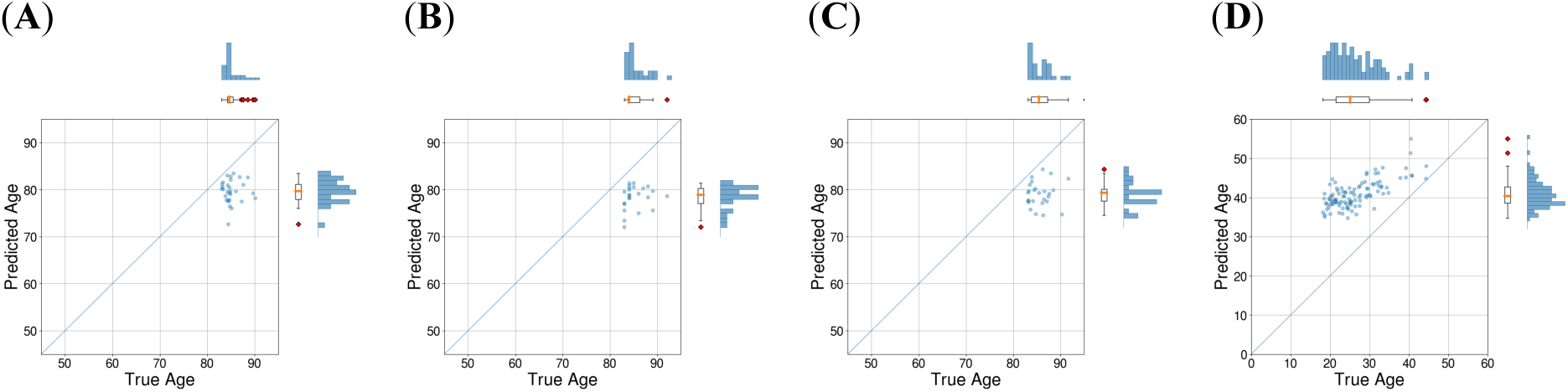
Extrapolation ability of Model #3. (**A**) Predicted brain age vs chronological age for subjects older than 81 in ADNI, (**B**) Predicted brain age vs chronological age for subjects older than 81 in AIBL, (**C**) Predicted brain age vs chronological age for subjects older than 81 in OASIS3, (**D**) Predicted brain age vs chronological age for subjects younger than 45 in MR-ART.

**Table 4.** Trained models’ extrapolation ability vs interpolation ability. Performance comparison of our model when predicting the chronological age of subjects younger than 45 and older than 83 (out of training age range), versus subjects older than 45 and younger than 83 (within training age range). Mean Absolute Error (MAE) measured in years. Prep: Our Preprocessing Pipeline, Aug: Data Augmentation, Drop: Input Dropout (20%).

| Dataset | ADNI |  | AIBL |  | OASIS3 |  | MR-ART |  |
| --- | --- | --- | --- | --- | --- | --- | --- | --- |
| Independent Testing Data Out of Training Age Range |  |  |  |  |  |  |  |  |
| n | 30 |  | 24 |  | 27 |  | 107 |  |
| [Min, Max] | [83.0, 90.1] |  | [83.0, 92.0] |  | [83.1, 95.3] |  | [18.2, 44.3] |  |
| Mean ± Std | 85.1 ± 1.80 |  | 85.2 ± 2.38 |  | 86.0 ± 2.87 |  | 26.4 ± 6.15 |  |
| Model / Metric | MAE<br>[Corrected] | R2<br>(p-value) | MAE<br>[Corrected] | R2<br>(p-value) | MAE<br>[Corrected] | R2<br>(p-value) | MAE<br>[Corrected] | R2<br>(p-value) |
| Model #1<br>SFCN-reg + Prep | 8.04<br>[6.17] | 0.02<br>(NS) | 8.56<br>[6.75] | 0.08<br>(NS) | 8.34<br>[5.92] | 0.01<br>(NS) | 18.48<br>[15.66] | 0.52<br>(p << 0.01) |
| Model #2<br>SFCN-reg + Prep + Aug | 7.41<br>[4.81] | 0.10<br>(NS) | 7.73<br>[5.28] | 0.01<br>(NS) | 8.34<br>[5.92] | 0.01<br>(NS) | 17.41<br>[13.14] | 0.55<br>(p << 0.01) |
| Model #3<br>SFCN-reg + Prep + Aug + Drop | 7.47<br>[5.45] | 0.00<br>(NS) | 8.67<br>[6.81] | 0.03<br>(NS) | 8.85<br>[6.92] | 0.01<br>(NS) | 17.31<br>[14.55] | 0.49<br>(p << 0.01) |
| Independent Testing Data within the Training Age Range |  |  |  |  |  |  |  |  |
| n | 250 |  | 220 |  | 586 |  | 12 |  |
| [Min, Max] | [59.9, 82.7] |  | [60.0, 82.0] |  | [46.0, 82.9] |  | [45.2, 74.8] |  |
| Mean ± Std | 73.5 ± 4.71 |  | 72.3 ± 5.42 |  | 67.1 ± 7.59 |  | 60.5 ± 8.85 |  |
| Model / Metric | MAE<br>[Corrected] | R2<br>(p-value) | MAE<br>[Corrected] | R2<br>(p-value) | MAE<br>[Corrected] | R2<br>(p-value) | MAE<br>[Corrected] | R2<br>(p-value) |
| Model #1<br>SFCN-reg + Prep | 2.85<br>[2.57] | 0.61<br>(p << 0.01) | 3.35<br>[3.04] | 0.61<br>(p << 0.01) | 2.70<br>[2.64] | 0.84<br>(p << 0.01) | 2.64<br>[2.27] | 0.93<br>(p << 0.01) |
| Model #2<br>SFCN-reg + Prep + Aug | 2.66<br>[2.63] | 0.65<br>(p << 0.01) | 3.41<br>[3.28] | 0.56<br>(p << 0.01) | 2.77<br>[2.89] | 0.81<br>(p << 0.01) | 2.18<br>[1.86] | 0.92<br>(p << 0.01) |
| Model #3<br>SFCN-reg + Prep + Aug + Drop | 2.84<br>[2.72] | 0.60<br>(p << 0.01) | 3.50<br>[3.20] | 0.56<br>(p << 0.01) | 2.86<br>[2.91] | 0.80<br>(p << 0.01) | 2.62<br>[2.59] | 0.89<br>(p << 0.01) |

Given the limitation of our models to extrapolate, in the following tests we consider only those subjects aged between 45 to 83 years old (the training age range) in order not to confound the MAE with extrapolation error.

### Test #3: Scan-rescan stability

An often-overlooked aspect of model assessment is the stability of its predictions. A reliable model should consistently generate similar results when presented with input from the same subject. The availability of multiple scans from the same subject within the same session in some datasets allows us to explore possible discrepancies in the model’s predictions when assessing the brain age of an individual from two distinct scans captured within hours during a single session. This evaluation of prediction stability is important to ensure the model’s reliability and its ability to produce consistent results under varying conditions, ultimately contributing to the overall robustness of the predictive framework. Combining ADNI and OASIS3, there are almost 500 subjects with two scans. We presented both scans to the models and calculated the absolute difference between the predicted ages. The mean absolute differences (MAD) and R2 score of first prediction and second prediction are reported in Table 5 and scatter plots are shown in Fig. 5 for each model and dataset (Note that we compare *differences* between two estimates and not the error between an estimate and the true age, hence the use here of MAD in the place of MAE).

**Figure 5.**
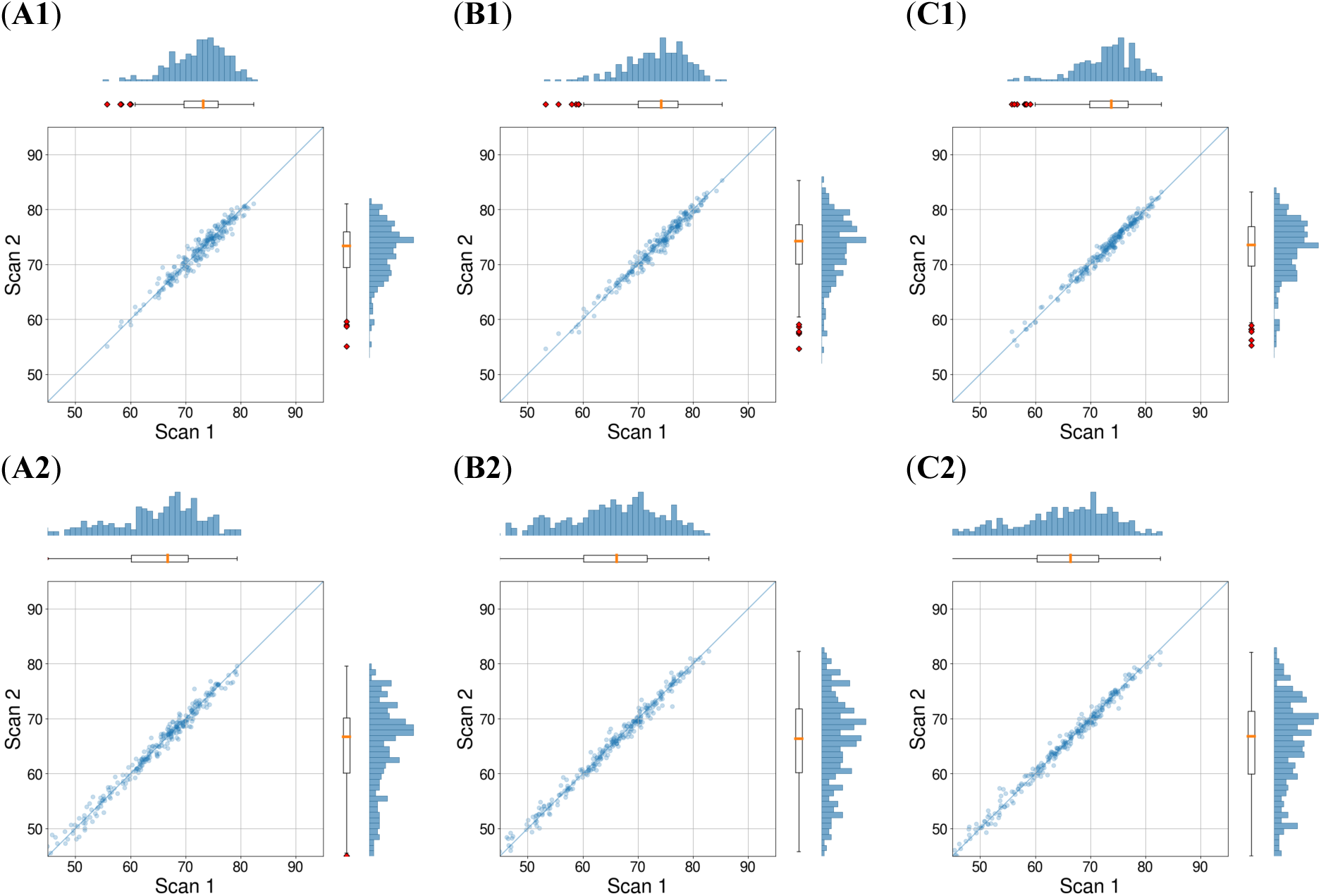
Scan-rescan stability of trained models. (**A1**) Model #1 - scan #2 predicted brain age vs scan #1 predicted brain age in ADNI, (**A2**) Model #1 - scan #2 predicted brain age vs scan #1 predicted brain age in OASIS3, (**B1**) Model #2 - scan #2 predicted brain age vs scan #1 predicted brain age in ADNI, (**B2**) Model #2 - scan #2 predicted brain age vs scan #1 predicted brain age in OASIS3, (**C1**) Model #3 - scan #2 predicted brain age vs scan #1 predicted brain age in ADNI, (**C2**) Model #3 - scan #2 predicted brain age vs scan #1 predicted brain age in OASIS3.

**Table 5.**
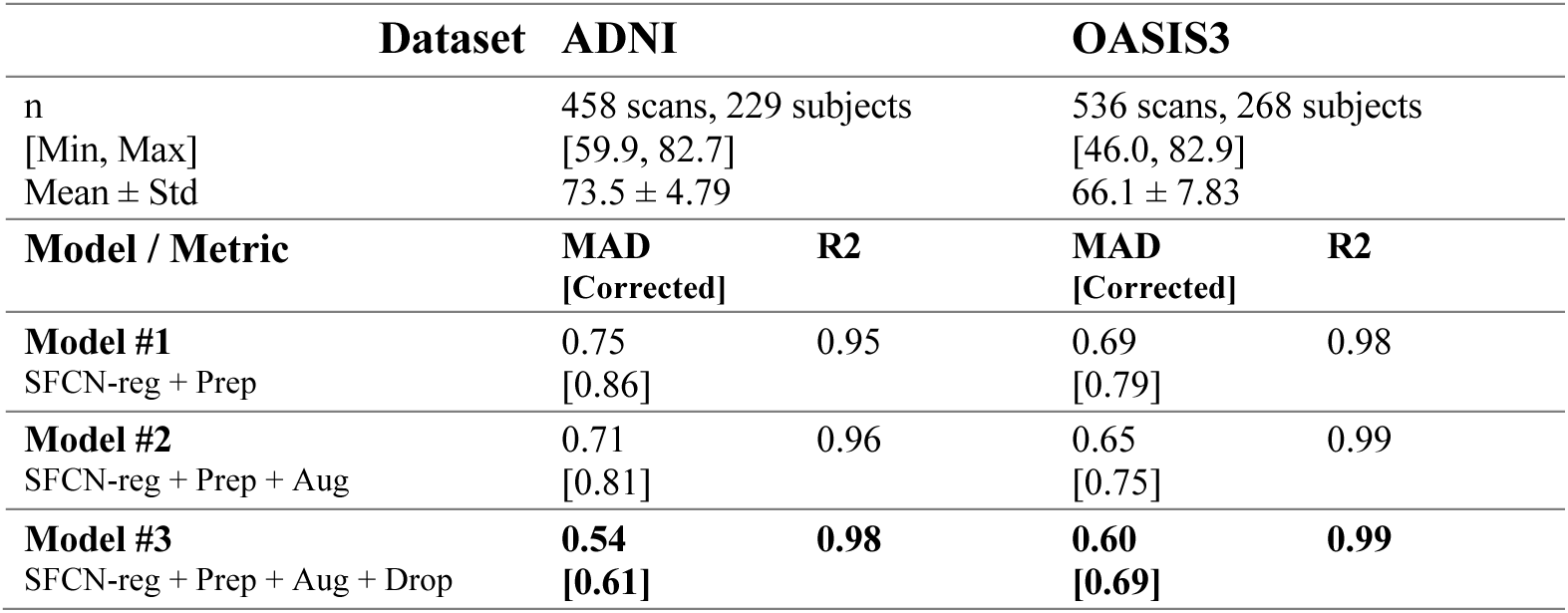
Comparison of the stability of our three models when presented with two different scans of the same subject during the same session. Mean Absolute Difference (MAD) measured in years [after bias correction]. Prep: Our Preprocessing Pipeline, Aug: data Augmentation, Drop: Input Dropout (20%).

Correlation plots (after bias correction) are shown in Fig 5. While all three models performed well with R2 scores of 0.95 or greater on different datasets, Model #3 trained with augmentation and input dropout was superior in comparison to the other two (up to 29.1% better than Model #1 and up to 24.7% better than Model #2 in terms of MAD). Likewise, Model #2 that was equipped with extensive data augmentations during the training, performed up to 5.8% better in comparison with Model #1 in terms of MAD.

### Test #4: Sensitivity to registration errors

As the performance of the trained models heavily relies on the data preprocessing pipeline, it becomes imperative to assess its sensitivity to potential processing issues such as registration errors. To conduct this evaluation, we selected a random subset of 50 scans from different test sets (e.g., from n=1885 UKBB held-out data, n=250 ADNI, n=220 AIBL, and n=586 OASIS3) and subjected them to 50 random affine transformations each. The goal was to quantify the model’s response to registration errors, with a focus on understanding the impact of such variations on predictions. The random transformation parameters included uniformly chosen scale parameters ranging from 0.98 to 1.02 independently for each axis, uniformly chosen rotation parameters spanning from −2 to 2 degrees independently for each axis, and uniformly chosen translation parameters covering −2 mm to 2 mm independently for each axis. Then, the predicted brain age for the perturbed input was compared with that for the unchanged input.

As shown in Table 6, Model #3 outperforms Model #1 and Model #2 in terms of Signed Difference (SD), indicating a smaller prediction bias. For different datasets, there is an absolute bias of 0.09 years on average for Model #3, while the average of absolute bias for Model #1 and Model #2 is 0.37 and 0.31 years respectively. Moreover, Absolute Difference (AD) improved by up to 9.3% from Model #1 to Model #2 and improved by up to 11.1% from Model #2 to Model #3 on average.

**Table 6.**
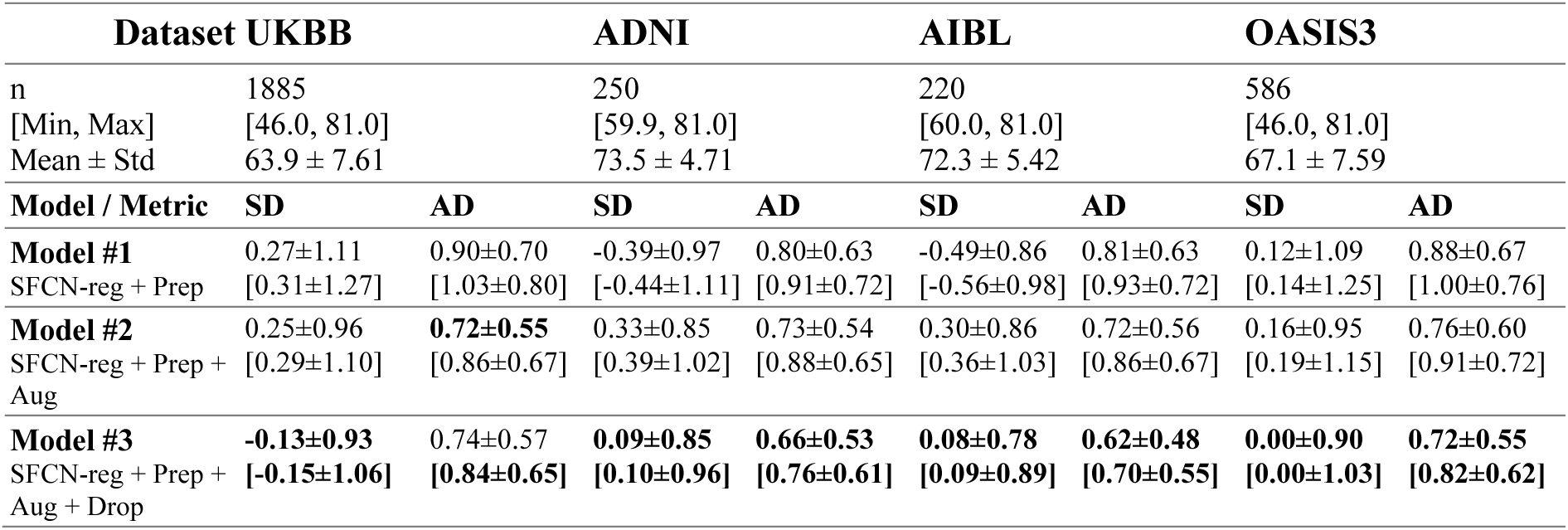
Comparison of the sensitivity of our three models to registration errors. Signed Difference (SD) and Absolute Difference (AD) measured in years [after bias correction]. Prep: Our Preprocessing Pipeline, Aug: Data Augmentation, Drop: Input Dropout (20%).

### Test #5: Sensitivity to motion artifacts

As shown in [35], brain age prediction can be affected by varying levels of motion artifacts. Therefore, it is important to evaluate the impact of motion on model outputs and minimize it as much as possible. To assess the impact of motion artifacts on our model, we compared the predicted brain age in the presence and absence of movement on the MR-ART dataset. The more robust the model, the less disparity there is between model outputs. According to test #2, our model does not perform well when it comes to subjects younger or older than the training age range. Hence, we exclude those subjects when performing this test to solely assess the impact of movement and omit the impact of instability our model encounters when facing younger or older subjects. MR-ART rated the level of movement with scores from 1 to 3. Score 1 indicates no or minimal movement, while 2 and 3 are associated with more severe motion artifacts. We compared the prediction for scans with a score of 1 and scans with a score greater than 1 for each subject and then calculated the MAD and R2 score for the entire population. Model #2, which was trained on images with motion augmentation, performs better than Model #1, showing an improved R2 score by 14.1% (from 0.78 to 0.89) and MAD by 20.3% (from 4.58 to 3.65 years). Model #3 also performs better compared to Model #1 (R2 score: 0.90 and MAD: 4.04 years), but worse than Model #2 in terms of MAD. The results are also reported in Table 7.

**Table 7.** Comparison of the stability of our three models when presented with two different scans of the same subject with different levels of movements during the same session. Mean Absolute Difference (MAD) measured in years [after bias correction]. Prep: Our Preprocessing Pipeline, Aug: Data Augmentation, Drop: Input Dropout (20%).

| Dataset | MR-ART |  |
| --- | --- | --- |
| n | 28 |  |
| [Min, Max] | [45.2, 74.8] |  |
| Mean ± Std | 61.0 ± 8.10 | |
| Model / Metric | MAD | R2 |
| **Model #1** | 4.00 | 0.78 |
| SFCN-reg + Prep | [4.58] |  |
| **Model #2** | **3.03** | 0.89 |
| SFCN-reg + Prep + Aug | **[3.65]** |  |
| **Model #3** | 3.54 | **0.90** |
| SFCN-reg + Prep + Aug + Drop | [4.04] |  |

### Interpretability of the Model

To analyze the contribution of various features to the output, we used Grad-CAM [36]. In summary, Grad-CAM examines the gradient of the output with respect to different features. While Grad-CAM is typically utilized to examine final layers, having our input image registered to a template allows us to apply Grad-CAM to the input layer. Fig. 6 displays averaged Grad-CAM maps of the Model #3 for different age groups (incrementing by 5 years). According to Grad-CAM maps, the model typically looks at the ventricle area and cortical areas. For younger subjects, the model focus is more towards the ventricle area, but for older subjects’ cortical areas are important as well. This is in alignment with our previous knowledge about brain aging, confirming that the model is paying attention to biologically plausible features.

**Figure 6.**
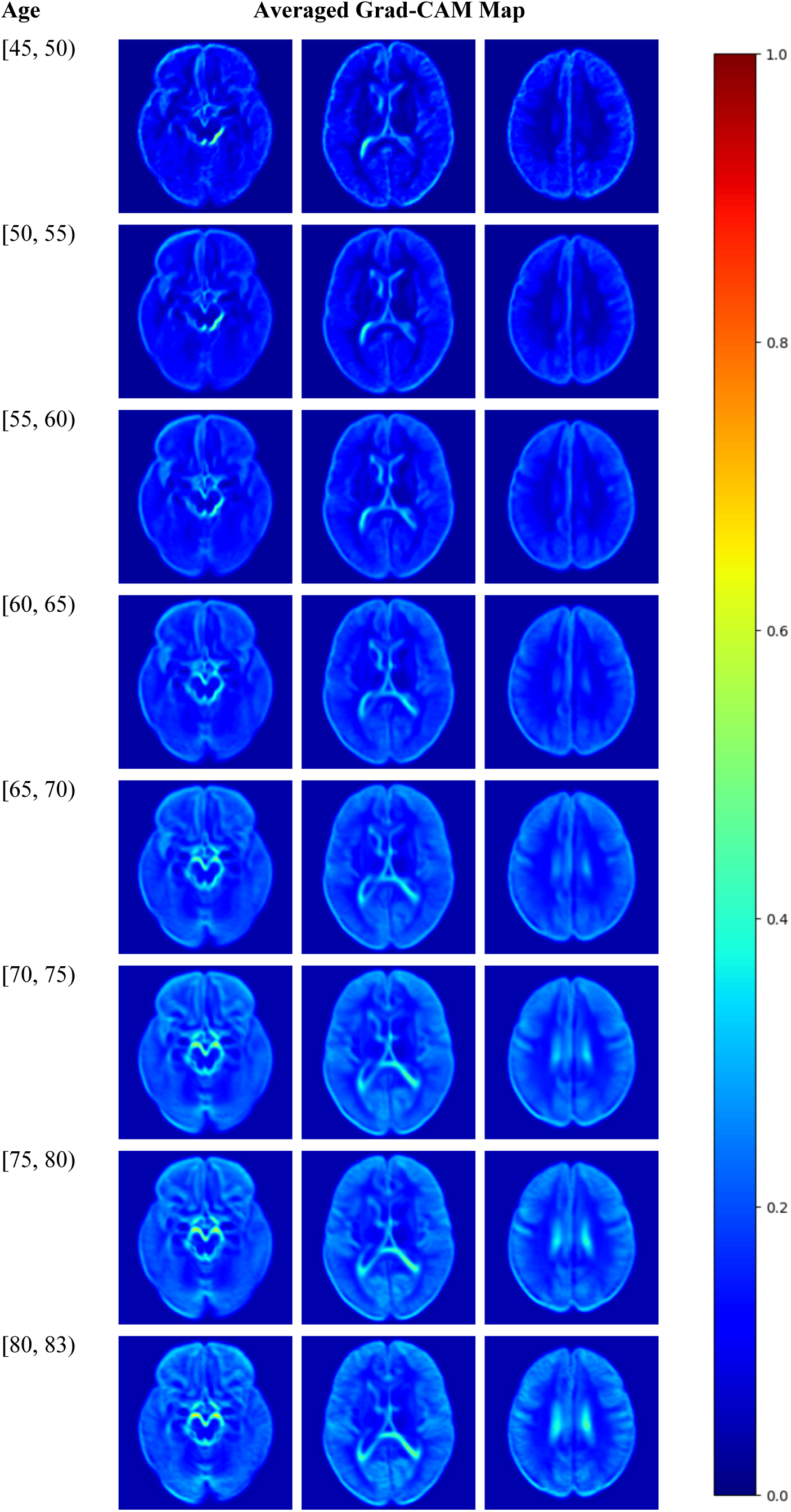
Averaged Grad-CAM Map for different age groups.

## Discussion

Many deep learning models have previously been trained and deployed to estimate an individual’s brain age out of T1w MR brain scans alone. Those models were reported to be more accurate than traditional methods. However, in our review of the literature we found that only a few of them were validated on out-of-distribution independent data that was not used for training [9], [17], [18], [19]. Moreover, we have not found any methods that assess the generalization and robustness of the trained model on more than one external dataset, thus most methods do not address the fact that other independent samples may have other distribution characteristics due to different factors such as scanner properties, preprocessing pipeline, inclusion/exclusion criteria, etc. Here, we trained our model on almost 40K images from the UKBB, a very large epidemiological sample, representative of the UK population, and tested it not only on a held-out within-distribution subset of the UKBB data, but also on out-of-distribution independent data from ADNI, AIBL, OASIS3 and MR-ART datasets. Furthermore, we conducted various tests to assess the reliability and robustness of our deep learning brain age prediction model and applied the following approaches to see if we could increase the reliability and robustness of the model: 1-implementing a more comprehensive preprocessing pipeline (Model #1), 2-incorporating additional data augmentation during training (Model #2), and 3-applying model regularization through an input dropout (Model #3).

By employing the aforementioned techniques and training the same model (SFCN-reg) with the same hyperparameters as used by Leonardsen et al. [17], we observed an improvement in brain age prediction generalization gap on all four independent datasets: ADNI, AIBL, OASIS3, and MR-ART. When applied to the ADNI dataset, Leonardson [17] reported an increase of 1.43 years in MAE, comparing within domain to out of domain testing. Here, we have an increase in MAE of only 0.86 years for Model #1, 0.65 years for Model #2, 0.70 years for Model #3, indicating that the proposed preprocessing, data augmentation and input input dropout lead to better generalization. One should note that Lee et al. [18] report only a slight increase (from 3.48 to 3.51 years) for within domain to out of domain testing, but at 3.5 years their MAE is more than 40% greater than that reported here. More specifically, the MAE of the SFCN-reg architecture of Leonardsen [17] of 5.25 years on the ADNI external validation set was reduced to 2.96 years with the more extensive preprocessing proposed here. This represents a 43.6% decrease in brain age predictive error. Smaller, but still important improvements (21.8%) were found for the AIBL dataset.

Adding more extensive data augmentation did not further decrease the average MAE in the external test data. However, data augmentation improves the robustness of the model as seen in the scan-rescan experiment (Test 3), the registration sensitivity experiment (Test 4) and the MR-artifact experiment (Test 5).

Although preprocessing images can improve the generalization, it also introduces potential sources of error into the model, such as errors in masking, non-uniformity correction, and registration, it is even more necessary to assess the reliability of the model with regards to these kinds of errors.

In the scan-rescan test, all models performed well with a minimum R2 score of 0.95, with Model #3 emerging as the top performer. Absolute difference decreases from 0.86 years with preprocessing, to 0.81 years with preprocessing and data augmentation, and to 0.61 years for preprocessing, data augmentation and input dropout for ADNI. However, considering that Model #3 is slightly less accurate in predicting brain ages of subjects from external independent datasets, the improvement in the scan-rescan test may be attributed to the increased bias of this model. Similar improvements were found with OASIS3.

In the registration sensitivity experiment, we showed that the bias is reduced almost to zero with input dropout, and the variance decreases as data augmentation and dropout are added.

In the sensitivity to motion artifacts experiment, it is evident that brain age predicted by our model is sensitive to motion. This is not surprising as many other preprocessing pipelines are sensitive to motion artifacts as well [37]. Moqaddam et al. [35] have shown that brain age prediction using conventional methods is also sensitive to motion and needs correction. However, as shown in Table 7, data augmentation helped us to decrease the disparity between two scans by different levels of movements by ∼20%.

We investigated how well our model could extrapolate when presented with samples older than those used in training. Our results (Table 4 and Fig. 4) revealed a pronounced floor/ceiling effect, indicating that none of our three models extrapolate effectively, particularly for older subjects, possibly due to their more atrophied brains or data distributions which are unfamiliar to the model. Neither augmentation nor regularization improved the extrapolation results. Despite this limitation and training our model on a smaller dataset with a narrower age range, testing it on samples from ADNI, AIBL, and OASIS3 (without age restrictions) resulting in enhanced results compared to the literature, enabling us to infer that better image preprocessing can enhance the model’s generalization ability more effectively than introducing additional training data via data augmentation or regularizing the model by input dropout. This improvement has also been reported by Dular et al. [19]. It is noteworthy to mention that our model outperforms those trained by Dular et al. [19], however direct comparison is challenging since we trained our model on 10 times more data and tested it on several datasets. We also used more steps in our preprocessing pipeline. Additionally, Dular et al. [19] employed non-linear registration in their pipeline, which may eliminate certain information related to brain atrophy that might be useful in brain age prediction.

Fig. 6 shows the anatomical features used to drive the brain age estimate. The earliest age range shows that it starts with information around the brain stem and lateral ventricles. As age progresses, more information from ventricles and medial temporal region is used, followed by even more information from the ventricles and the cortex. This follows our knowledge about brain aging, indicating reliability of features extracted by our model.

Our study is not without limitations. As demonstrated by the extrapolation test, our models are unable to predict the brain age of subjects younger than 45 or older than 81. To better evaluate these deep learning models, it appears necessary to train them on a broader, more inclusive age range, similar to what discussed by Kopal et al. [38], [39]. These deep models lack interpretability on their own, and their inability to extrapolate raises even more questions about their interpretability. Our models rely on a preprocessing pipeline, requiring preprocessing for every image introduced to the network. Our preprocessing pipeline is optimized to execute in less than 15 minutes per image on an average desktop computer. However, it is still significantly slower than pipelines that rely solely on deep learning architectures processing raw images. When registration is used as part of the preprocessing pipeline, the same stereotaxic MRI template is employed to prevent bias in the model. While the template we used is suitable for adults, it may not be optimal for young children or for older subjects with significant brain atrophy. Finally, the architecture utilized is well-established in brain age prediction literature but does not incorporate several more recently developed techniques, such as attention mechanisms, which may enhance accuracy.

In conclusion, we have demonstrated that application of a sophisticated image preprocessing pipeline, extensive data augmentation during training, and regularization through input dropout can significantly improve the accuracy and robustness of brain age prediction in healthy datasets. Evaluation experiments in the left-out within distribution UKBB data and in the ADNI, AIBL, OASIS3 and MR-ART independent datasets show good performance and strong out-of-distribution generalizability, low scan-rescan variability, and robustness to registration errors. On top of that, the Grad-CAM images show that anatomically appropriate features are used to estimate brain age. In the future, we will apply this model to patient cohorts with cognitive decline and or neurodegeneration.

